# Florigen revisited: proteins of the FT/CETS/PEBP/PKIP/YbhB family may be the enzymes of small molecule metabolism

**DOI:** 10.1101/2021.04.16.440192

**Authors:** Olga Tsoy, Arcady Mushegian

## Abstract

Flowering signals are sensed in plant leaves and transmitted to the shoot apical meristems, where the formation of flowers is initiated. Searches for a diffusible hormone-like signaling entity (“florigen”) went on for many decades, until in the 1990s a product of plant gene *FT* was identified as the key component of florigen, based on genetic evidence and protein localization studies. Sequence homologs of FT protein are found throughout prokaryotes and eukaryotes; some eukaryotic family members appear to bind phospholipids or interact with the components of the signal transduction cascades. We studied molecular features of the FT homologs in prokaryotes and analyzed their genome context, to find tentative evidence connecting the bacterial family members with small molecule metabolism, often involving sugar- or ribonucleoside-containing substrates. Most FT homologs share a constellation of five charged residues, three of which, i.e., two histidines and an aspartic acid, circumfere the rim of a well-defined cavity on the protein surface. We argue that this conserved feature is more likely to be an enzymatic active center than a catalytically inactive ligand-binding site. We propose that most of FT-related proteins are enzymes operating on small diffusible molecules, which may constitute an overlooked essential ingredient of the florigen signal.

## Introduction

Flower production in plants occurs in response to the environmental cues – most importantly, changes in the day length. The role of photoperiodicity in all living forms from bacteria to higher eukaryotes is well established, and it has been shown in numerous studies that the perception of the photoperiodic signal in plants occurs primarily in the leaves. Flowers, however, are formed mostly by shoot apical meristems (SAM), which are typically shielded from direct light, and the mechanisms which convey the flowering signal from leaves to SAMs have been a matter of speculation and investigation for most of the 20^th^ century.

Plant physiologists have established, by the 1930s, that angiosperms generally fall in three categories, i.e., long-day plants, in which blooming is turned on by day lengthening / night shortening; short-day plants, which initiate blooming upon day shortening / night lengthening; and day-neutral plants, which bloom in response to the cues other than day length (Garner and Allard, 1920; Knott, 1934). Experiments on flowering in partially shaded plants and in grafts between light-induced and uninduced plants suggested the existence of a flower-promoting diffusible substance; the idea may have been first proposed by Julius Sachs (Sachs, 1865), but was consolidated in the modern form by M. Chailakhyan (1902-1991), who proposed the term “florigen” as the name for the flower-inducing chemical entity (Chailakhyan, 1936). The early research on the photoperiodic signal sensing in leaves is reviewed in (Zeevaart, 2006; Kobayashi and Weigel, 2007), and an account of Chailakhyan’s remarkable life and research work can be found in (Romanov, 2012).

The transfer of a flowering signal from leaves to the shoot apex has been studied over the years in many species of flowering plants, and general similarity of its properties to those of small molecules was noted; for example, the florigen fraction appeared to move in phloem at the speed comparable to that of other plant assimilates (King et al., 1968). It has been established also that grafts and extracts of induced plants could transfer flowering ability not only in the uninduced vegetatively developing plants of the same species, but sometimes also in other species or genera, suggesting the evolutionary conservation of the signaling pathways (Zeevaart, 1976).

Despite general acceptance of the idea of florigen as a conserved hormone-like substance, the attempts at the isolation and chemical characterization of the responsible entity did not succeed for several decades. Reviewing the state of the affairs in 1976, J.A.D. Zeewart listed the factors that have been tested unsuccessfully for the ability to induce plant flowering; the list of failed candidates included sugars, amino acids, sterols, gibberellins, salicylic acid, ethylene, cytokinins, the photosynthetic capacity, initiation of protein synthesis, and others (Zeevaart, 1976). Several hypotheses have been put forward to explain the difficulties of identifying florigen: perhaps it was not one molecule but several distinct hormones or metabolites, acting jointly in a specific succession, or as a mix with specific ratio of components; or, possibly, the identity of florigen was masked by simultaneous presence of flowering inhibitors in the same samples (Lang et al., 1977); or, maybe, flowering was caused by propagation of electric, rather than chemical, signals between leaves and meristems, such as plasma membrane potential (Penel et al., 1985).

Despite all the work to test these and other hypotheses, there was little progress in biochemical characterization of florigen until early 1990s, when the search for a flower-inducing activity started employing the tools of molecular biology. For example, in what may have been the last published study by Chailakhyan, a radiolabeled protein band of ~27 kDa was observed in the induced, but not in naïve, leaves of *Rudbeckia*, followed by accumulation of the similar-sized band in the shoot apex tissues (Milyaeva et al., 1991); the work is largely unavailable to the Western reader because of the temporary interruption of translation and indexing of Russian-language journals upon the collapse of the Soviet Union. Around the same time, it has been shown (Takeba et al., 1990) that aqueous extracts of *Lemna, Pharbitis* and *Brassica* contained a flower-inducing fraction dominated by a 20-30 kDa protein band; the authors noted, however, that the florigen activity was resistant to proteinase K digestion that removed the protein, perhaps suggesting a role for an associated small molecule.

Nearly simultaneously, the results of genetic screens for *A.thaliana* mutants with late flowering phenotypes were published (Koornneef et al., 1991). This was a watershed moment in the studies of the molecular determinants of flowering initiation, and the important discoveries that ensued in the following three decades have been reviewed extensively (Andrés and Coupland, 2012; Lifschitz et al., 2014; Kinoshita and Richter, 2020). The state of the knowledge may be summarized as follows.

Flowering in Arabidopsis is LD-dependent, and is enabled through the circadian clock-controlled transcriptional co-regulator *CONSTANS (CO)* and its target *FLOWERING LOCUS T*. The protein product of the latter gene has emerged as the integrator of the environmental inputs, relaying these signals into the gene regulatory network that controls flowering. FT protein is produced in the phloem companion cells of the leaves, enters the phloem sieve elements and is transported from the leaves to the base of shoot apical meristem, where it has been detected experimentally. Considerable evidence exists that the *FT* gene is not expressed in SAM above its basal portion, and that the FT protein may move from cell to cell in plants. This is a key set of properties expected of florigen. A strong genetic and transgenic evidence suggests that FT is required for activating floral meristem identity genes and flowering time controllers, such as MADS-box transcription factors SUPPRESSOR OF OVEREXPRESSION OF CONSTANS 1 (SOC1), FRUITFULL (FUL/AGL8) and AGL24, and ultimately the master regulator of flower development LEAFY (LFY). Orthologs of FT, such as SINGLE FLOWER TRUSS (SFT) in tomato, HEADING DATE 3a (HD3a) in rice, and their counterparts in many other species (sometimes called CETS proteins in the plant context, after the names of several better-studied representatives from *Antirrhinum, Arabidopsis* and *Lycopersicon*) share many of these properties with FT, with some variation in the precise wiring of the transcriptional networks. The paralogs of *FT*, found in varying numbers in all examined angiosperm species, also tend to be involved in flowering control, acting either synergistically or antagonistically with *FT*, though additional or exclusive roles in other aspects of plant development have been shown for a subset of them (Abelenda et al., 2019; Ahn et al., 2006; Conti and Bradley, 2007; Danilevskaya et al., 2011; Jin et al., 2021; Krieger et al., 2010; Pin and Nilsson, 2012; Pnueli et al., 2001; Tsuji et al., 2011).

Despite all effort, FT protein has not been detected in phloem-free tips of SAM in any plant, only in the basal cells of the meristem zones. Protein localization studies at a single-cell resolution in vivo remain challenging, and it is not clear whether FT actually reaches the cells where the meristem identity genes are expressed. The argument that FT does just that – as opposed, for example, to activating an additional low molecular weight messenger – has been made in the literature, but the evidence is indirect, showing for example that some engineered fusions of FT to green fluorescent protein-based reporters are retained in the phloem at the bottom of the SAM zone, and in those cases they do not complementation the *ft* recessive mutants (Corbesier et al., 2007). A study in rice (Tamaki et al., 2007) has made a more direct claim that Hd3a, the protein ortholog of FT in rice, does reach the tip of the SAM to induce flowering. However, their Fig. 3, cited as the key supporting evidence, does not seem to show the protein reporter activity at the tip, where it is supposed to be. Therefore it remains possible that FT is but one component of the flower induction signal, and that a small molecule, possibly activated by or acting synergistically with FT, is also involved (Lifschitz and Eshed, 2006; Andrés and Coupland, 2012).

Another gene of Arabidopsis involved in flowering control, *FD*, has been identified as a recessive suppressor of *FT* – the ability of overexpressed FT to induce precocious flowering is impaired in plants with the lowered production of FD (Abe et al., 2005). The *FD* gene encodes a bZIP-type transcription factor, which interacts with the FT protein in yeast two-hybrid system, in the bi-molecular fluorescence complementation assays in vivo, and apparently in the affinity-purified protein complexes (Abe et al., 2005; Wigge, 2005). In rice, the FT ortholog HD3a does not interact with the bZIP factors directly, but the two proteins co-precipitate in a tripartite complex that also includes a scaffolding 14-3-3 protein (Taoka et al., 2011). These interaction data are the major part of the evidence on the basis of which FT has been assigned the molecular function of a transcriptional co-activator, proposed to form a putative multisubinit complex that binds to the regulatory regions and controls the expression of the flowering identity genes. A corroboration of these protein interaction experiments is provided by the analysis of gene expression and ChIP-Seq data, which reveal that tagged FT binds, apparently mostly in the FD-dependent manner, near many of the genes involved in flowering control (Zhu et al., 2021).

Analysis of the amino acid sequence has shown that FT protein belongs to a widely conserved sequence family, members of which are encoded by genomes of many bacteria, archaea and nearly all eukaryotes. The founding member of the family, isolated from bovine brain, is a cytoplasmic protein that can bind in vitro to many low molecular weight compounds of different chemical structure, including certain phospholipids (Bernier and Jollès, 1984). The name phosphatidylethanolamine-binding protein (PEBP) became attached to the family, though functional relevance of phosphatidylethanolamine binding has not been demonstrated for any homolog of this protein in any species. The genes or protein products of FT/PEBP family turn up in many genetic screens and binding assays. This has resulted in a variety of putative properties assigned to these proteins, including, in addition to phospholipid binding in plants and animals, also inhibition of carboxypeptidase Y and regulation of Ras GTPase in yeast, inhibition of Raf-1 kinase in mammals (hence an alternative name of the protein family, RKIP), suppression of transepithelial migration of the mammalian host neutrophiles by a secreted, plasmid-encoded PEBP homolog from uropathogenic *E.coli* (Lau et al., 2014), and an uncharacterized role in the modification of polyketide chains in *Streptomyces* (Li et al., 2008, 2009).

Structural studies of crystallized FT/PEBP-like proteins from diverse species of bacteria and eukaryotes have revealed a distinct spatial fold, with a structural core dominated by two beta-sheets and superficially resembling an immunoglobulin-like arrangement (entry 11.1.7 in the ECOD database (Cheng et al., 2015)). A prominent structural feature of this family is an evolutionary conserved cavity on the surface of the molecule. The cavity sometimes accommodates anions included in the crystallization media, including the phosphoryl moiety of the co-crystallized phosphoethanolamine, but is neither hydrophobic nor large enough to bind lipids; on the other hand, several sites suitable for binding hydrophobic ligands have been inferred on the molecule by in silico studies, but those sites tend to be located in the least conserved regions of the molecule (Roderick et al., 2002; Ahn et al., 2006; Shemon et al., 2010; Rudnitskaya et al., 2012; Guo et al., 2018; Nakamura et al., 2019). Curiously, the site of interaction between FT and FD in *A.thaliana* has not been structurally characterized.

In this work, we argue that the patterns of sequence and structure conservation in the family, as well as the genomic context of the FT/PEBP proteins in prokaryotes, are best compatible with the idea that these proteins are enzymes involved in production, attachment or removal of low molecular weight ligands, and that some of such small molecules could play a role in induction of flowering in angiosperms. Thus, rather than consisting only of the FT protein itself, florigen’s active moiety may comprise a low molecular weight product of an enzymatic activity of FT.

## Results

### Sequence conservation in the FT homologs throughout the Tree of Life suggests shared ancestry and common molecular function

We have collected the homologs of plant FT proteins by the PSI-BLAST searches of the NCBI NR database, focusing mostly on completely sequenced genomes. We also consulted the NCBI COG resource that annotates conserved orthologous genes in bacteria and archaea. In the following, unless specified otherwise, we refer to the prokaryotic orthologs from this family as YbhB proteins, after the name of the chromosomal gene in *E.coli*, and use some of the FT/CETS/PEBP/PKIP name aliases for eukaryotic homologs. Complete genomes of the unicellular and multicellular eukaryotes tend to encode at least one, or commonly more than one, FT homolog; a rare exception are some parasitic eukaryotes with reduced genomes, such as microsporidia, which do not appear to have genes from this family. In bacteria and archaea, the distribution of YbhB orthologs is more complex. YbhB genes (NCBI COG1881) are found in almost all major lineages of bacteria and archaea. Among the clades of archaea, only methanogenic *Euryarchaeota* appear to lack the YbhB homologs, and among bacteria, only the species in phylae *Firmicutes*, *Mollicutes* and in the order *Spirochaetales* tend to be YbhB-free. In most other clades of bacteria and archaea, between 30 and 80% of all species encode YbhB homologs. All told, there are 782 copies of YbhB family proteins found in 594 bacterial and archaeal genomes out of the 1309 genomes in the 2020 release of the COG database. YbhB homologs are also encoded by the genomes of many giant DNA viruses from the *Megaviricetes* class and from some related lineages.

We produced a multiple sequence alignment of the representative proteins from many of these clades and inferred a phylogenetic tree of these sequences. The alignment is shown in Figure 1, and the tree in the Newick format is available as the Supplemental Data Set 1. Many of the branches in the phylogenetic tree are long, and the position of a few clades is not well-resolved. The partitioning of prokaryotic sequences in the tree, however, shows close correspondence to the phylogenetic positions of the species in which they are found; the statistically supported clades generally do not mix bacterial and archaeal representatives, and it is only occasionally that a sequence from a phylogenetically distant bacterium is found within a group of more closely related homologs, as in the case of one sequence from Gram-negative proteobacterium *Helicobater pylori* nested inside a group of YbhB-like sequences from Gram-positive bacteria. All this indicates that the evolution of the YbhB family in prokaryotes must have included the episodes of rapid evolution and differential gene loss, while the horizontal gene transfer events may have been relatively rare in this family.

**Figure 1.**
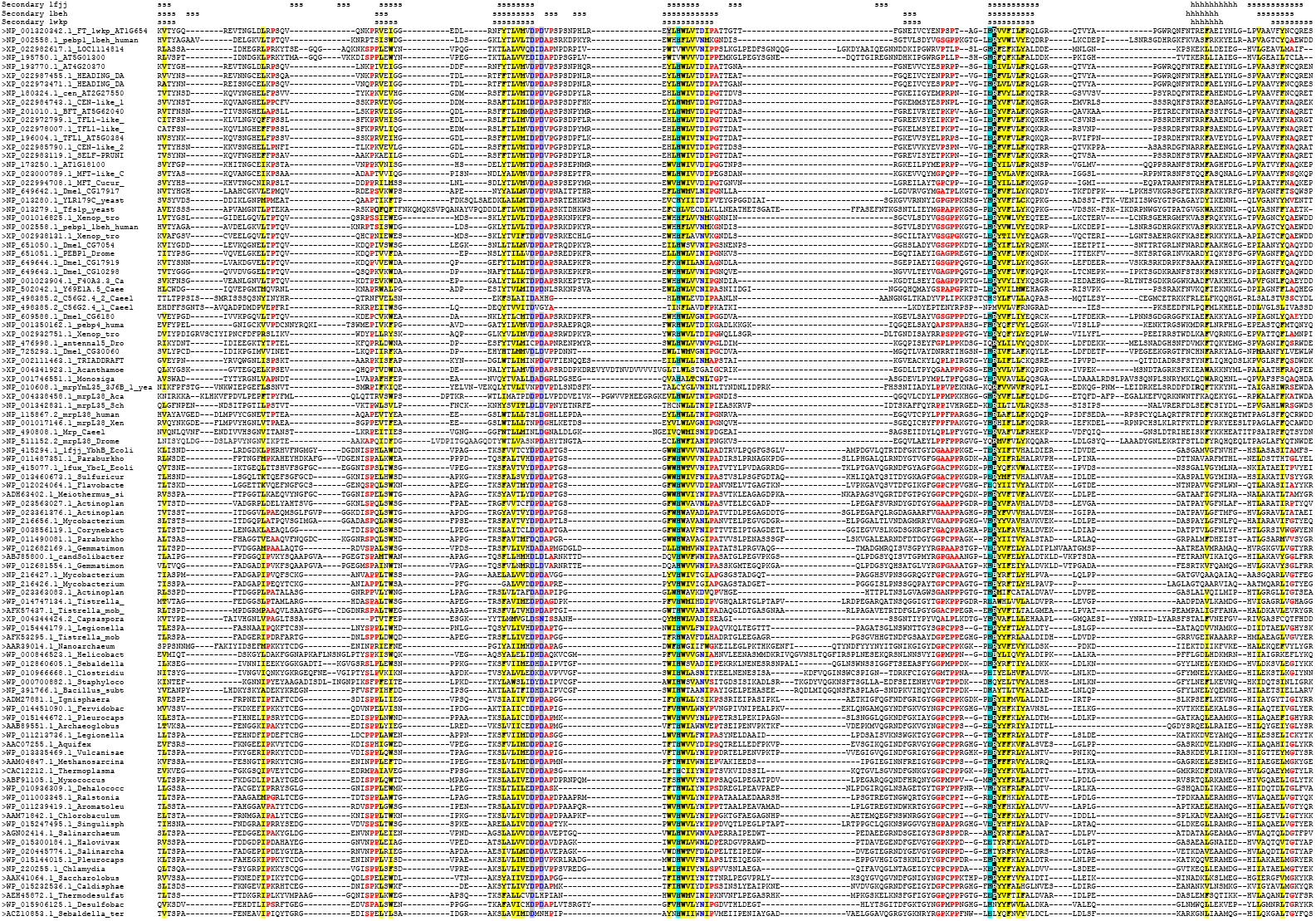
Multiple sequence alignment of select members of FT/CETS/PEBP/PKIP/YbhB family. Sequence identifiers in GenBank or PDB are shown before each sequence. In the Secondary structure lines above the alignment, s stands for a beta-strand and h stands for an alpha-helix. Conserved hydrophobic residues (I, L, M, V, F, Y, W) are indicated by yellow shading, conserved small-side-chain or turn/kink-prone residues (A, G, S, P) are indicated by bold red type, and the conserved constellation of charged residues discussed in the text are marked as follows: gray-shaded blue type, two conserved aspartates; cyan shading, two conserved histidines; and black-shaded white type, conserved arginine.

The eukaryotic sequences are separated from bacterial and archaeal sequences by a long internal branch in the unrooted tree. Within eukaryotes, nearly all plant FT/CETS homologs segregate as one clade, and fungal/metazoan homologs represent another assembly, occasionally intermingled with the representatives of unicellular eukaryotes; evolution and specialization of FT homologs in different plant lineages has been extensively discussed, e.g., (Klintenäs et al., 2012; Karlgren et al., 2011; Jin et al., 2021). Within fungal and metazoan proteins, a distinct clade includes FT/PEBP homologs detected recently in an unexpected biological context, namely as the constituents of mitochondrial ribosomal large subunit, where they are known as mitochondrial ribosomal proteins (Mrp) L35/L38. These FT/PEBP/YbhB homologs have distinct N-terminal sequence extensions, found in one gene product within each completely sequenced animal, fungal and protist genome (many of animal and fungal genomes also include additional, extension-free homologs). This domain is not included in the alignment in Figure 1; none of the plant FT homologs appear to contain such a region.

Taken together, these data suggest the ubiquity of the FT/CETS/PEBP/PKIP/YbhB-like proteins throughout the evolution of life and a single origin of all eukaryotic homologs from a prokaryotic ancestor. It is likely also that all or most of those proteins share an ancient conserved molecular function, reflected in the nearly-universal sequence conservation of several key amino acid residues and the common structural scaffold of the entire protein family (Figure 1, and see the next section).

### Structural analysis of PEBP/PKIP/FT/YbhB proteins supports the hypothesis of an enzymatic function

A common biochemical function of the entire PEBP/YbhB protein family is further suggested by mapping of the conserved and variable sequence elements of the family onto the available three-dimensional structures (Figures 1 and 2). There are five nearly-invariant polar amino acid residues in the alignment – two aspartic acids located at the C-terminus of strand 3, two histidines found at the N-termini of strands 4 and 6, and an arginine that adjoins the second conserved histidine. Substitutions in these positions in prokaryotes are rare, and multiple replacements are seen mostly in the eukaryotic mitochondrial ribosomal proteins L35/L38, which is the clade that is most likely to have acquired a new molecular function during evolution. There are also several highly conserved residues, in particular prolines, in the loops around these signature amino acids (Figure 1). The five most conserved residues are also found close to each other in the spatial structures of the proteins (Figure 2).

**Figure 2:**
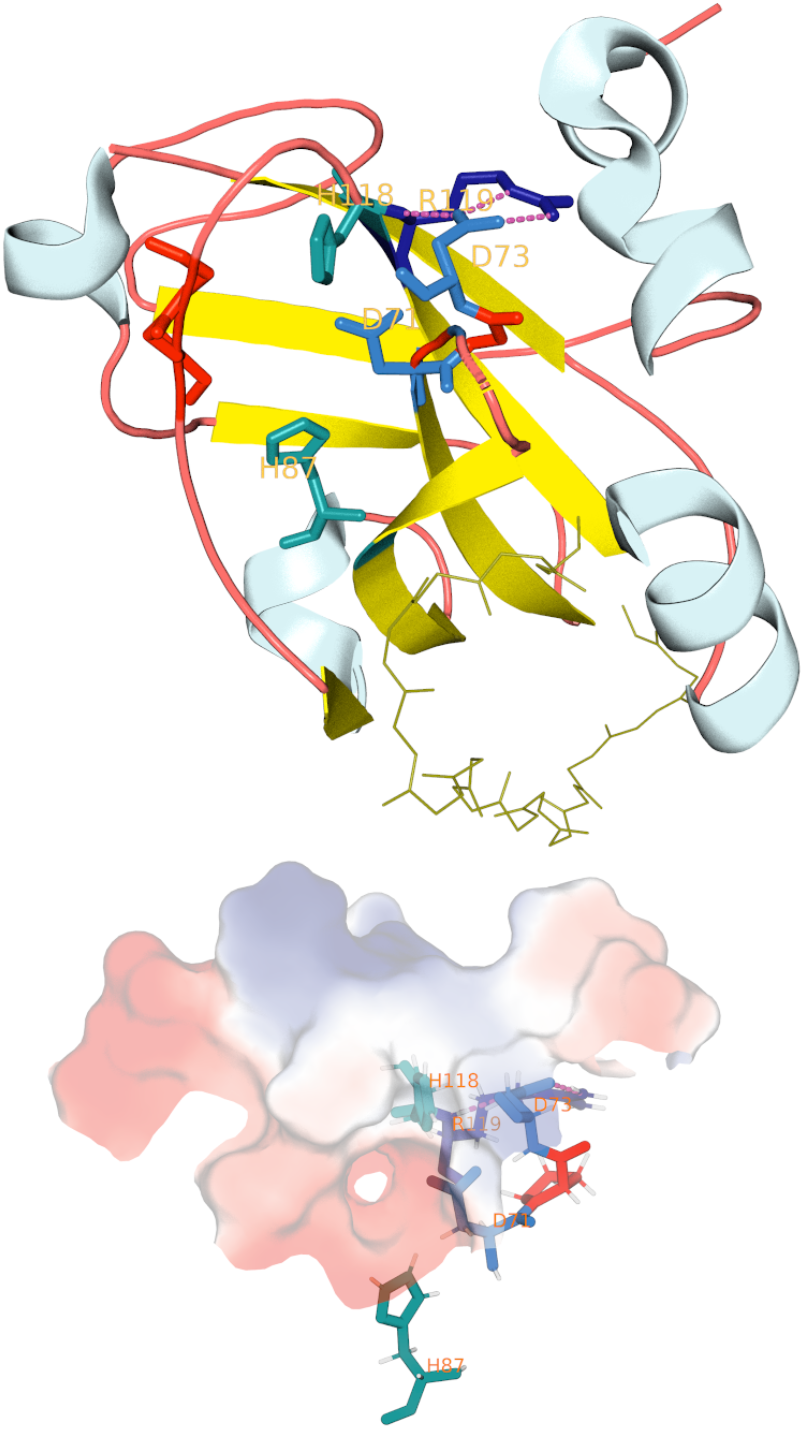
Spatial organization of the conserved residues forming putative catalytic center in FT protein from Arabidopsis tlialiana (PDB ID 1wkp). Top pane, strands are rendered in yellow, helices in light blue, and conserved residues involved in forming the putative enzyme active center are labeled and colored as follows: blue, two conserved aspartates; cyan, two conserved histidines; dark blue, conserved arginine; red, frequently conserved prolines. The loop between strands 3 and 4 is reduced to a short broken wire to improve the visibility of the putative active center, and the loop that is a major determinant of the antagonistic activity between FT and it paralog TFL1 (Ahn et al., 2006) is rendered as sticks. Bottom pane, the putative enzyme active center rendered as a surface. Rough calculation of the surface electrostatic properties in the vacuum were performed using PyMOL generate → vacuum electrostatics function; the shades of red indicate negative charges, and the shades of blue indicate positive charges. The conserved residues involved in forming the putative enzyme active center are labeled and colored as in the top pane.

Both histidines, the first of the two aspartates, and the proline-rich loop before the strand 6 form the rim of a hydrophilic hole, which is the largest cavity conserved in all structurally characterized proteins in the family. The second of the conserved aspartates forms forms three hydrogen bonds with the side chain of the conserved arginine, constituting a conserved “shoulder” on the surface of the molecule (Figure 2).

The polar cavity on the surface of all PEBP/PKIP/FT/YbhB protein molecules has been interpreted as a site that binds a ligand containing a polar group, such as phosphate; many of the described or suspected interactions of eukaryotic PEBP/PKIP/FT with phospholipids and phosphoproteins have been rationalized in this way. Recently, a report that FT from Arabidopsis may bind phosphatidylcholine (PC) more strongly and more specifically than other lipids has prompted the analysis of high-resolution structures of FT crystallized in the presence of PC. Unexpectedly, in the crystals obtained under various conditions no electron density for PC could be detected, and computational docking suggested four potential binding sites for PC, all located far from the conserved polar cavity (Nakamura et al., 2014a, 2014b). It is also notable that the electrostatic calculations suggest that the overall charge of the cavity, at least in vacuum, is strongly negative, making it perhaps not so conducive for direct interactions with phosphate, whereas the charge on the “shoulder” may be more positive (Figure 2B). On the whole, the attempts to relate the known biological properties of the FT homologs to their sequence and structure still do not provide a satisfactory explanation of the role of the most prominent, conserved elements of the family.

We suggest that these signature elements of sequence and structure are indicative of an enzymatic function of the FT/YbhB homologs. The search of Mechanism and Catalytic Site Atlas (M-CSA) resource, which contains annotated information about the amino acid determinants of catalysis in conserved enzyme families, identifies 21 families of enzymes that have two His, two Asp and one Arg in their active centers. These families represent all 7 of the top-level Enzyme Classification classes of activities, i.e., oxidoreductases, transferases, hydrolases, lyases, isomerases, ligases and translocases. If the selection criteria are relaxed to only two histidines and one aspartic acid surrounding the hydrophilic hole, the search retrieves 105 families, corresponding to almost 11% of the 964 entries in the database. The particular three-dimensional configuration of these residues in PEBP/PKIP/FT/YbhB proteins appears to be quite unique, as judged by the analysis with the PINTS program, which compares similar spatial arrangements of key residues in non-homologous proteins (Stark and Russell, 2003). Nonetheless, it is clear that a combination of two histidines, two aspartates and a lysine has been repeatedly utilized in evolution to build enzymatic active centers, enabling catalytic conversions of different kinds.

These observations are in sharp contrast to what is known about sequence conservation in non-catalytic ligand-binding protein domains. A collection of such domains was extracted from the PFAM database, using keywords “ligand+bind” and “phosphate-binding”, resulting in 1044 conserved domains. Removal of the clearly annotated enzymatic domains that bind phosphate in their active centers, such as the protein kinases or ATPases, and selection of a non-redundant set of sequence families, produces 651 putative non-catalytic ligand-binding domains. Examination of the sequence profiles of these families suggests that the majority of the conserved positions in these domains tend to be occupied by non-polar residues, although conserved charged residues, such as lysine, histidine, arginine, glutamate or aspartate, are also found occasionally. We recorded the identity of all charged residues that were characterized by the information content of more than 50% of the maximum possible value for their position, and found only 25 domains that had more than two conserved charged residues; none of those domains simultaneously had two conserved histidines and a conserved aspartate (Supplemental File 2). Thus, the known conserved ligand-binding domains do not appear to use the combination of two histidines and an aspartic acid residue for their non-catalytic interactions with small molecules, suggesting again that the conserved feature in PEBP/FT/YbhB proteins is more likely to be a part of the catalytic center than to serve solely as a binding interface.

### Genomic context of FT homologs in bacteria and yeast suggests connections with small molecule metabolism

We took advantage of the broad taxonomic distribution of the YbhB homologs and tried to infer the putative functional linkages for these gene products on the basis of their genome context (Table 1). One kind of a linkage between proteins may be revealed when two protein domains exist as separate open reading frames in some species, but are fused into multidomain proteins in others; such translational fusions, especially those that are evolutionary conserved, are enriched in proteins that work in concert with one another (Huynen et al., 2000; Suhre and Claverie, 2004). The YbhB homologs in bacteria and archaea are most commonly encoded as stand-alone open reading frames and form domain fusions infrequently. In actinobacteria, however, they are found in the C-terminal portions of longer proteins, which also contain the modules implicated in carbohydrate metabolism. For example, protein WP_014179790.1 in *Streptomyces sp*. consists of the N-terminal pectate lyase-like carbohydrate-binding module (domain ID in NCBI CDD database cd04082), followed by the FN3 repeat region (cd00063), a putative glucose/sorbosone dehydrogenase (GSDH) region with predicted beta-propeller structure (cl268170), and finally the C-terminal YbhB homology domain. This theme is partially preserved, albeit with domain rearrangement, in some species of evolutionarily distant proteobacteria, where the order of domains is GSDH-YbhB, or occasionally GSDH-FN3-YbhB; examples include WP_014747134.1 in an alphaproteobacterium *Tistrella* and proteins in gammaproteobacteria, such as WP_096298086.1in *Luteimonas*, PYD93448.1 in *Pseudomonas syringae* pv. *pisi*, or WP_145513070.1 in *Xantomonas perforans*. GSDH enzymes utilize quinone cofactors to convert hexoses and their derivatives into a variety of products in bacteria (Miyazaki et al., 2006).

**Table 1.**
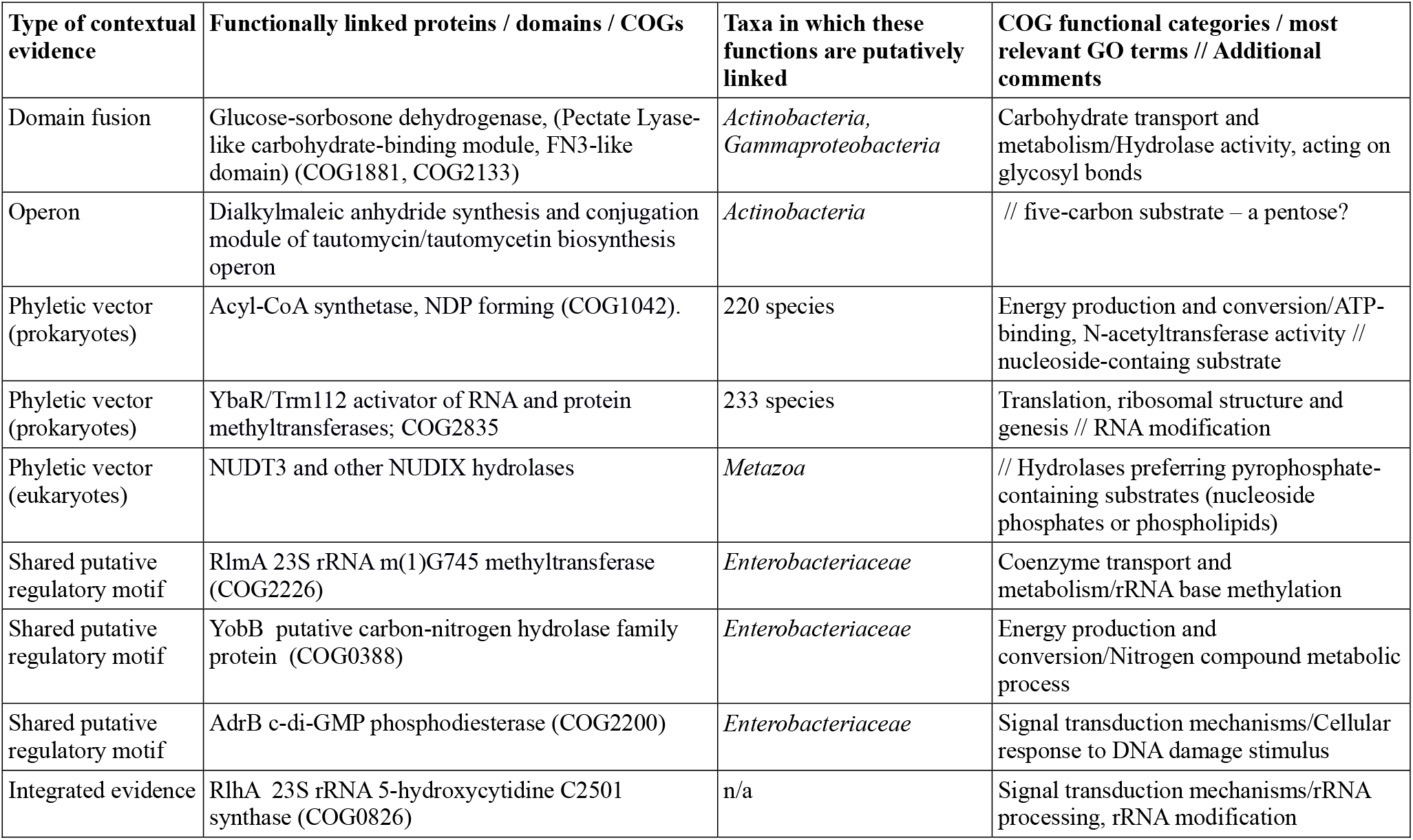
Genome-context information suggesting functional links between YbhB homologs and various enzymes of biosynthesis and salvage of small molecules. See text for a more detailed characterization of each putative functional link.

We then surveyed the databases of conserved genomic neighborhoods and scanned the literature for the evidence about the conservation of FT/YbhB chromosomal neighbors in different genomes. As with the protein fusions, the FT/YbhB family genes are not commonly found within predicted operons or have conserved positional associations with other ORFs. In two species of *Streptomyces*, however, YbhB homologs are located within the large (20 genes) operons responsible for biosynthesis of polyketide-based small molecules, i.e., tautomycin in *S.spiroverticillatus* and tautomycetin in *S.griseochromogenes*. These polyketides are of pharmacological interest, because they are potent inhibitors of mammalian protein phosphatases. The YbhB family genes, called, respectively, *ttnL* and *ttmL*, together with the adjacent seven open reading frames, are required to manufacture a rare dialkylmaleic anhydride moiety of tautomycin and tautomycetin and to conjugate it to the polyketide backbone, produced by the remaining genes in the same operon. Dialkylmaleic anhydride is required for the activity of the mature product, and is made *de novo* from propionate and an unidentified C5 compound, through the succession of the steps that are incompletely understood (Li et al., 2008, 2009).

Another approach of connecting genes into functionally related groups is to analyze their phyletic vectors, i.e., the representations of the presences and absences of their orthologous genes in different genomes (Glazko and Mushegian, 2004; Glazko et al., 2006). We have searched the space of phyletic vectors associated with the 2014 release of NCBI COGs, using the vector of YbhB (COG1881) as the query. After retaining the matching vectors with comparable cardinality (i.e., the genes present in 300-400 species, to avoid spurious matching to almost-ubiquitous proteins), we found two COGs that were most often co-inherited by the same genomes as COG1881. One of those, COG1042, is annotated as Acyl-CoA synthetase (NDP forming); it is found in 358 species, 220 of which encode both COG1881 and COG1042. The enzyme, best studied in hyperthermophilic archaea and protists, is involved in the substrate-level phosphorylation, by the equation acetyl-CoA + ADP + Pi ⇌ acetate + ATP + CoA (Musfeldt and Schönheit, 2002). The roles of the bacterial homologs is less clear, as some of them appear to be catalytically inactive and possibly play auxiliary roles in the acylation and deacylation of proteins (Thao and Escalante-Semerena, 2012). The other gene with matching phyletic pattern, COG2835, annotated as “uncharacterized conserved protein YbaR, Trm112 family”, is found in 390 genomes, 233 of which also encode COG1881. YbaR/Trm112 family encodes activators of several methyltransferases involved in modification of rRNA, tRNA and peptide release factors (van Tran et al., 2018)

One more way to predict functional linkages between gene products is to examine their putative regulatory regions, which in bacteria tend to be located in the intergenic regions, usually to the 5’ ends of the open reading frames or operons they regulate. We performed such analysis of YbhB homologs in several bacterial taxa and detected a conserved site upstream of the YbhB open reading frame in the family *Enterobacteriaceae*. Unlike most known binding sites for the specialized transcription factors, this site lacks palindromic structure and has a consensus sequence TACACTT. Scanning 41 *Enterobacteriaceae* species from 14 genera with a probabilistic model of the site identifies additional intergenic regions where the variants of these sites occur. (Table 1, and Supplemental File 3). Three genes that were found to contain a highly similar conserved upstream site in 16-17 species, representing 8 genera are *rlmA* (23S rRNA m(1)G745 methyltransferase), *yobB* (putative carbon-nitrogen hydrolase family protein), and *adrB* (c-di-GMP phosphodiesterase). Sites matching this consensus in *E.coli* have been noticed before and predicted to represent a modified form of the canonical −10 element sequence TATAATT (Huerta and Collado-Vides, 2003); it is located in an appropriate position relative to the ORFs starts, but the regions in which we found this upstream element do not appear to have the recognizable −35 sequence nearby. The significance of the shared TACACTT element thus remains unclear, though a common regulation mechanism for genes that share this site is a possibility.

Finally, we have consulted several curated databases of pre-computed functional linkages. PEBP/FT/YbhB homologs are not well-represented among the modules reported by those databases, but two observations were of interest. The FunCoup database, which uses naïve Bayesian approach to integrate the information from 10 different genomic and proteomic measurement spaces (Persson et al., 2021), shows a small network of interacting proteins in *E.coli*, which includes two chaperones, i.e., a heat shock 70-family DnaK and a protease Lon, as well as YbhB and YbcL. Interactions with chaperones, which are frequently observed in the protein interaction data, may reflect cellular proteostasis needs and a broad clientele of many chaperone systems, but are often not particularly informative about the protein function. The connection to the other protein in the group, YbcL, may be more revealing: that protein, recently renamed RlhA, appears to be a component of the 5-hydroxycytidine synthase enzymatic complex involved in modification of 23S rRNA (Kimura et al., 2017). And the analysis of phyletic patterns across many eukaryotes, collected in PhyloGene database (Sadreyev et al., 2015) has revealed one highly-scoring match co-inherited with human PEBP1 gene, i.e., NUDT3, encoding an enzyme from the Nudix class. Nudix proteins are hydrolases noted for the affinity to the pyrophosphate moieties in their substrates, often lipid, nucleoside or oligonucleotide derivatives (McLennan, 2006).

## Discussion

> “Circumstantial evidence is a very tricky thing,” answered Holmes thoughtfully. “It may seem to point very straight to one thing, but if you shift your own point of view a little, you may find it pointing in an equally uncompromising manner to something entirely different.”
>
> — Arthur Conan Doyle. *The Boscombe Valley Mystery*.

We present the computational evidence indicating that the molecular function of the FT proteins in flowering, as well as the biochemical roles of the FT/CETS/PEBP/PKIP/YbhB homologs, could have been misunderstood: the proteins from this family might be the enzymes involved in the biochemical transformation of small molecules, either instead of, or in addition to, their postulated role of being stoichiometric subunits within plant transcription complexes.

Much of the analysis reported here comes from the examination of prokaryotic genome sequences. Obviously, studies in archaea and bacteria cannot be expected to validate the role of FT in the transcriptional control of flowering in plants, as prokaryotes lack most of the plant signal transduction systems and downstream effectors of flowering. Instead, we asked two different questions, i.e., “What can be deduced about the molecular functions of the FT orthologs in prokaryotes?” and “Is there evidence that these functions may be conserved in eukaryotes, including plants?”

The first of these questions can be tentatively answered by taking into account the genomic context of FT/YbhB homologs hinting at their possible functional connections to enzymes involved in specific metabolic pathways. Though no single gene could be linked to YbhB by several orthogonal approaches, a trend emerges when these data are considered jointly (Table 1). The evidence appears to point towards functional linkages of YbhB to sugar and ribonucleotide/RNA modifications, implying that FT/YbhB may be involved in the metabolism of a monosaccharide such as ribose or another pentose, or perhaps their nucleoside-like derivative. Relatedly, FT proteins could be involved in phospholipid metabolism that proceeds through lipid conjugates with nucleotides; indeed, recent genetic evidence has suggested the involvement of phosphorylethanolamine cytidylyltransferase (*PECT1* gene product) in flowering in Arabidopsis plants (Susila et al., 2021). Finally, FT proteins could have a role in attaching or removing some low molecular weight moieties from larger molecules, such as proteins or nucleic acids; this could explain the association of FT and its homologs with nuclear proteins seen in plants and other eukaryotes.

The answer to the other question posed above, i.e., whether the putative enzymatic function could have been preserved through the evolution of the FT/CETS/PEBP/PKIP/YbhB family, appears to be clearly in the affirmative; the pattern of sequence conservation and spatial juxtaposition of the key charged residues in the family at a long phylogenetic span is striking (Figures 1 and 2), even though this constellation of conserved residues has not been definitely mapped to a specific biological function thus far. Bioinformatic analysis suggests that the conserved sets of two histidines and an aspartic acid clustered in space are frequent in the enzyme active centers but virtually never are found in the non-catalytic ligand-binding domains. Finally, broad occurrence of the protein family in bacteria, archaea, giant viruses and nearly all eukaryotes would not be unusual for an enzyme, especially one involved in metabolism or signaling, but would be a much rarer occurrence for transcriptional regulators, as they are typically not shared between bacteria and eukaryotes (Babu et al., 2004).

There is an ample precedent of utilization of sugars, nucleotides and products of RNA breakdown for plant hormone biosynthesis; one class of plant hormones, cytokinins, are purine derivatives that can be produced either by isoprenylation of adenosine phosphate or by tRNA degradation (Kakimoto, 2003), whereas another class, gibberellins, are synthesized by transforming the pentose skeleton generated in the 1-deoxy-D-xylulose 5-phosphate pathway (Salazar-Cerezo et al., 2018). Recent studies has significantly expanded the repertoire of linear and cyclic oligonucleotides that serve as essential signaling messengers in bacteria and animals (Burroughs et al., 2015; Burroughs and Aravind, 2020), again suggesting that some nucleoside derivatives with regulatory properties may remain undiscovered in plants.

A critic might point out that the bioinformatic evidence of the enzymatic function of FT and their homologs presented in this manuscript, let alone of the proposed involvement of FT in specific biochemical pathways, is too circumstantial – some would say, speculative – and thus far has not been corroborated by the wet-laboratory experiments. However, exactly the same can be said about the experimental support for the role of FT as a transcriptional co-activator – a hypothesis that has not been corroborated by the computational evidence and is quite unmoored from the facts of comparative genomics. In the spirit of considering the entire corpus of the available data, and using reciprocal illumination from different classes of evidence to improve the precision of scientific hypotheses (Iyer et al., 2001; Baker et al., 2014; Carr et al., 2015), it seems timely and urgent to start testing the hypothesis of a metabolic role of the FT homologs in various organisms. Comparison of metabolic profiles in the wild-type organisms and in FT knock-down mutants could be a first step in determining the identity of the substrates and products of the putative enzymatic activity that we predict to be conserved in most members of this protein family.

## Methods

Sequence homologs were collected using the PSI-BLAST program (Altschul et al., 1997) with standard settings, run either against the NCBI NR protein sequence database or the databases restricted by taxon (e.g., viruses). Nucleotide and amino acid sequences were aligned using the program MUSCLE v. 3.8.31 (Edgar, 2004). Distribution of the YbhB homologs in bacterial and archaeal genomes was studied using the web interface for the 2020 release of NCBI COG database (Galperin et al., 2021). The phylogenetic trees were constructed by PhyML program implemented in Phylogeny.fr server (Dereeper et al., 2008), with 1000 bootstrap replicates and approximate Likelihood Ratio Test. The three-dimensional structures of proteins were visualized and approximate, charge-smoothed electrostatic surface representations were generated using the open-source PyMOL environment (Schrödinger LLC; SciCrunch RRID SCR_000305), installed from source using homebrew on the MacOS High Sierra 10.13.6. Gene adjacency on the chromosomes and protein domain fusions were analyzed by visual inspection of the sequence records in GenBank. Phyletic vectors were analyzed using the psi-square program with default settings and vector similarity measured using Pearson correlation-based distance (Glazko et al., 2005); the NCBI COG database release of 2014 was used as the search space, because the digitized phyletic vector information was not available for the 2020 release at the time of writing. For promoter analysis, the complete bacterial genomes were downloaded from Ensembl Bacteria database (release 47) (Howe et al., 2020) in March 2020; if several strains were available for a species, one random strain has been chosen. The regulatory regions of homologous genes were aligned with MUSCLE and the most conservative regions were chosen to build a positional weight matrix (PWM) by the SignaX routine of the Genome Explorer program (Mironov et al., 2000). The genome scan was performed by Genome Explorer with the threshold set equal to the lowest score in the training set, i.e., in the set of *ybhB* homologous regions. The usage of specific amino acid residues by the enzymatic active centers was performed with the aid of Mechanism and Catalytic Site Atlas (M-CSA) online resource, using the Web interface to select the residues of interest (Ribeiro et al., 2018). The usage of specific amino acid residues by ligand-binding domains was studies by collecting all domains in PFAM 33.1 (El-Gebali et al., 2019) with keywords “ligand bind” or “phosphate-binding”. The known enzymatic domains were manually excluded based on their description, and for each remaining domain, the curated seed alignment has been downloaded from PFAM and used as an input for the Skylign server (Wheeler et al., 2014) to build HMM profiles and to get the information count for the residues of interest.

## Supplemental Data

Supplemental Data Set 1. Newick-formatted phylogenetic tree of FT/YbhB homologs in bacteria, archaea and eukaryotes.

Supplemental Data Set 2. List of ligand-binding domains in PFAM and their usage of conserved amino acid residues

Supplemental Data Set 3. Occurrences of the motif with consensus sequence TACACTT in *Enterobacteriaceae*.

## Acknowledgements and Authors’ Contributions

OT is supported by the German Federal Ministry of Education and Research (BMBF) within the framework of the e:Med research and funding concept (grant 01ZX1908A). AM is a program director at the National Science Foundation, an agency of the U.S. Government; his work was supported by the National Science Foundation’s Independent Research/Development and Long-Term Professional Development programs, but the statements and opinions expressed herein are made in the personal capacity and do not constitute the endorsement by NSF or the government of the United States.

AM designed the research; OT and AM performed research; OT and AM wrote the paper.

